# Olfactory coding in the bumble bee antennal lobe

**DOI:** 10.1101/2020.12.31.424929

**Authors:** Marcel Mertes, Julie Carcaud, Jean-Christophe Sandoz

## Abstract

Sociality is classified as one of the major transitions in evolution, with the largest number of eusocial species found in the insect order Hymenoptera, including the Apini (honey bees) and the Bombini (bumble bees). Bumble bees and honey bees not only differ in their social organization and foraging strategies, but comparative analyses of their genomes demonstrated that bumble bees have a slightly less diverse family of olfactory receptors than honeybees, suggesting that their olfactory abilities have adapted to different social and/or ecological conditions. However, unfortunately, no precise comparison of olfactory coding has been performed so far between honey bees and bumble bees, and little is known about the rules underlying olfactory coding in the bumble bee brain. In this study, we used *in vivo* calcium imaging to study olfactory coding of a panel of floral odorants in the antennal lobe (AL) of the bumble bee *Bombus terrestris*. Our results show that odorants evoke consistent neuronal activity in the bumble bee antennal lobe. Each odorant evokes a different glomerular activity pattern revealing this molecule’s chemical structure, i.e. its carbon chain length and functional group. Response intensity as well as odor-similarity relationships were highly correlated to those measured in honey bees. This study thus suggests that bumble bees, like honey bees, are equipped to respond to odorants according to their chemical features.

## Introduction

Sociality is classified as one of the major transitions in evolution, and animals often form social groups because the benefits (either direct or indirect) of grouping outweigh the costs of breeding independently (Bourke, 2011; Hamilton, 1964). The insect order Hymenoptera (including ants, bees and wasps) presents the largest number of eusocial species and up to 9 independent origins of eusociality (Rubenstein & Abbot, 2017). In bees, eusociality is notably found in the corbiculate bees (bees with concave “pollen baskets” on their hind legs) which include well-known eusocial taxa, the Apini (honey bees) and the Bombini (bumble bees). The ‘primitively eusocial’ bumble bees (*Bombus* spp.) share some traits with advanced eusocial species (like honey bees), but lack particular aspects that would qualify them as advanced eusocial organisms (Sadd et al., 2015). This intermediate position on the eusocial spectrum renders them a particularly interesting group for unraveling the mechanisms of social evolution.

Bumble bees and honey bees differ in many ways and show notable ecological differences (Sadd et al., 2015). Bumble bee colonies are annual and small, consisting of several hundred individuals, compared to the perennial honeybee colonies which contain many thousands of individuals. This means that the behavior of an individual bumble bee has a greater effect on the success of the colony than does the behavior of a single honey bee (Sherry & Strang, 2015). Division of labor in the colony also differs between honey bees and bumble bees. In honey bees, workers progress through various nest- and foraging tasks in an age-dependent fashion whereas in bumble bees, workers of all ages and sizes may perform nest or foraging duties (Jandt et al., 2009). With regards to foraging, larger bumble bees bring nectar to the colony at a higher rate but individuals of all sizes do forage (Spaethe & Weidenmüller, 2002), whereas in honey bees, nectar and pollen are collected only by a subgroup of older workers, the foragers. Social communication also differs. While both species use a number of pheromones within the nest, honey bees developed a unique symbolic communication system (the well-known dance language) to inform each other about the location of food sources (von Frisch, 1967). In the same context, bumble bees gather information from “excited runs” and pheromone signals provided by foragers returning to the nest (Dornhaus & Chittka, 2001; Dornhaus et al., 2009; Molet et al., 2009).

As mentioned upon above, chemosensation plays a major role in social interactions in insect societies, and is also critical for bees’ foraging success. Given the differences in social organization and foraging strategies existing between bumble bees and honey bees, we might expect important differences in how the two species process olfactory information. In insects, odorants are detected by olfactory receptors (ORs) carried by olfactory sensory neurons (OSNs) on the antennae (Fig. 1a). ORs belong to a multigenic family whose members are known to evolve quickly through complex patterns of gene birth and death (Ramdya & Benton, 2010). Comparative analyses of the genomes of a honey bee (*A. mellifera*) and two bumble bee species (*B. Impatiens* and *B. terrestris*) demonstrated that bumble bees have a slightly less diverse family of ORs than honeybees. Conversely, however, bumble bees possess an expanded repertoire of gustatory receptors (GR) compared to honey bees (Sadd et al., 2015), suggesting different priorities in the chemosensory systems of the two insects. Apart from absolute numbers, substantial differences are found between honey bees’ and bumble bees’ OR repertoires, with a limited number of ortholog genes. These observations suggest that their olfactory abilities have adapted to different social and/or ecological conditions. Unfortunately, no precise comparison of olfactory coding has been performed so far between honey bees and bumble bees.

**Figure 1:**
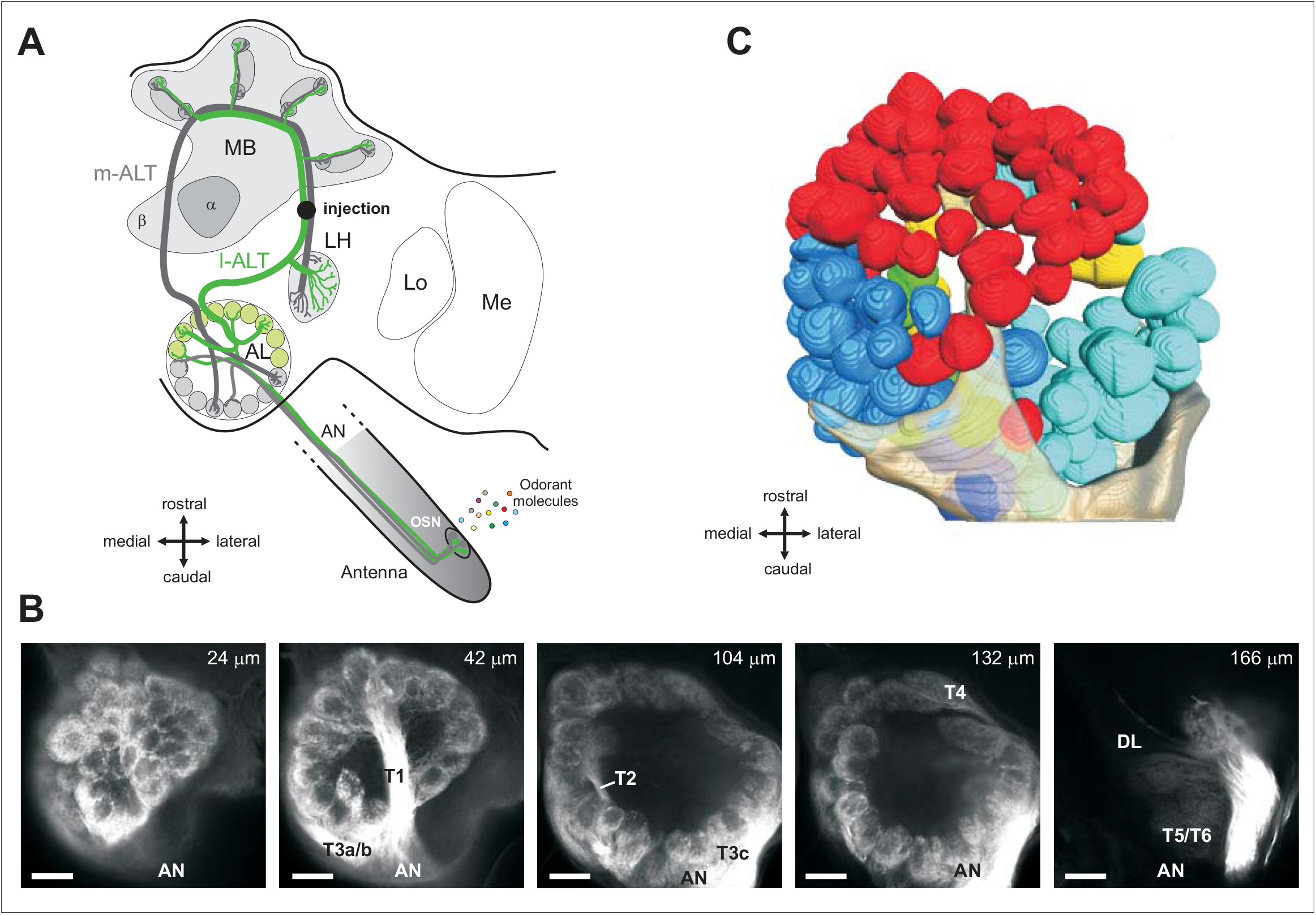
Anatomy of the bumble bee antennal lobe (AL). **(a)** Hymenopteran dual olfactory pathway (adapted from Carcaud et al. 2012). Odorant molecules are detected by olfactory sensory neurons (OSNs) on the antenna, which form the antennal nerve (AN) and send olfactory information to the primary olfactory center, the antennal lobe (AL). Then, projection neurons (PNs) convey information to higher-order centers, the mushroom bodies (MB) and the lateral horn (LH), using two main tracts, the l-ALT (lateral antennal-lobe tract, in green) and the m-ALT (medial antennal-lobe tract, in grey). PNs of the m-ALT and l-ALT project to distinct areas in the MB and in the LH. The black dot indicates the site of injection for calcium imaging. Lo: lobula, Me: medulla, α: α-lobe, β: β-lobe. **(b)** Confocal image sequence through a bumble bee antennal lobe (right lobe) obtained by massive anterograde antennal staining (using tetramethylrhodamine dextran). The scale bars indicate a length of 50 µm. The depth along the z-axis of the confocal images are indicated on the top right of each image. AN: antennal nerve. T1-T4: subdivisions of the antennal nerve in the AL. **(c)** Three-dimensional reconstruction of the 158 glomeruli in the antennal lobe presented in B. The glomeruli are colored depending on their input tracts. The numbers of glomeruli per input tracts are: T1 = 60 (red), T2 = 7 (green), T3a = 27 (medium blue), T3b = 16 (dark blue), T3c = 42 (turquoise), and T4 = 7 (yellow) glomeruli. The antennal nerve is shown in semi-transparent coloring.

Olfactory coding and processing have been intensively studied in the honey bee, a traditional animal model in neuroethology (Galizia et al., 2011; Giurfa, 2007; Menzel, 2012; Sandoz, 2011). In addition to extensive research on bees’ olfactory behaviors (Guerrieri et al., 2005; Nouvian et al., 2015; Vareschi, 1971), this insect’s olfactory pathways have been described in great details (Kirschner et al., 2006; Mobbs, 1982; Nishino et al., 2009) and physiological recordings like electrophysiology (Brill et al., 2013; Kropf & Rössler, 2018) and optical imaging (Jernigan et al., 2020; Joerges et al., 1997; Mota et al., 2013; Paoli et al., 2018; Szyszka et al., 2008) have unraveled the rules of odor coding. By contrast, less work has been devoted to the understanding of olfactory perception and learning in bumble bees, and most of this work used behavioral approaches (Laloi et al., 1999; Leonard & Masek, 2014; Riveros & Gronenberg, 2009; Sommerlandt et al., 2014). A number of studies have started to describe the anatomy of the bumble bee brain (Fonta & Masson, 1985; Mares et al., 2005; Paulk et al., 2008; Smith et al., 2016; Strube-Bloss et al., 2015). Its general architecture has been found to be highly similar to that of the honey bee, in particular with regards to the olfactory pathway (Fonta & Masson, 1985; Strube-Bloss et al., 2015). In both species, OSN project from the antenna to the primary olfactory center, the antennal lobe (AL), constituted of spherical anatomical and functional units, the glomeruli (∼165 in honey bees) (Hansson & Anton, 2000). Each glomerulus receives input from all the OSNs that express a given type of OR (Gao et al., 2000; Vosshall, 2000). Within the AL, local interneurons perform local computations (Meyer et al., 2013), and projection neurons (PNs) then convey processed information to higher-order centers, the mushroom bodies and the lateral horn. In honey bees and bumble bees, as in most Hymenoptera, the PNs are divided in two main tracts of uniglomerular neurons, the lateral antennal lobe tract (l-ALT) and the medial antennal lobe tract (m-ALT) (Abel et al., 2001; Kirschner et al., 2006; Strube-Bloss et al., 2015), with possibly different functions (Brill et al., 2013; Carcaud et al., 2015, 2018). Apart from the observation of a general similarity in the architecture of the olfactory pathway, functional studies of odor coding in bumble bees are scarce. In the 1980’s, two studies described bumble bees’ peripheral equipment in cuticular sensilla on the antennae and performed electroantennogram (EAG) recordings of their antenna, showing that it responds to a wide range of volatiles, including both floral and pheromonal odorants (Fonta & Masson, 1984, 1987). These approaches were used again later to show that sensillar equipment and olfactory sensitivity increase with worker size in bumble bees (Spaethe et al., 2007) as well as to study left-right asymetries (Anfora et al., 2011). With regards to neural odor coding, one study demonstrated the existence of glomerulus-size odor-induced oscillations in the bumble bee AL (Okada & Kanzaki, 2001). More recently, extracellular recordings of AL neurons showed reproducible responses to odorants, and observed a specific response pattern for a pheromonal compound compared to other odorants (Strube-Bloss et al., 2015). Apart from these findings, little is known about the rules underlying olfactory coding in the bumble bee brain.

In the present work, we used *in vivo* calcium imaging to study olfactory coding by l-ALT PNs in the AL of the bumble bee *Bombus terrestris*. First, to compare odor-coding rules in bumble bees and honey bees, we presented a panel of floral odorants previously used in studies on olfactory processing and perception in honey bees (Carcaud et al., 2018; Carcaud et al., 2012; Guerrieri et al., 2005). Our results show that odorants evoke consistent neuronal activity in the bumble bee antennal lobe. Each odorant evokes a different glomerular activity pattern depending on the molecules’ chemical structure, i.e. carbon chain length and functional group. Odor-similarity relationships in the bumble bee AL are highly correlated to those found in the honey bee AL.

## Results

### Anatomy of the bumble bee antennal lobe - olfactory sensory neuron innervation

Using fluorescent tracers, we performed mass staining of olfactory sensory neurons (OSNs) in the bumble bee *Bombus terrestris* (Fig. 1a). The tracers migrated along the antennal nerve until the OSNs’ axonal projections in the cortex (outer layer) of the glomeruli in the AL. As previously reported (Fonta & Masson, 1985), we found a similar arrangement of sensory tracts in the bumble bee antennal lobe as in the honey bee. The most prominent tract, T1, is easily identifiable, crossing the center of the antennal lobe from the antennal nerve caudally (Fig. 1b) to the most ventral and rostral part of the antennal lobe where it innervates many glomeruli. The T3 tract is also prominent, leaving the antennal nerve on the caudal side of the antennal lobe, propagating medially on its outskirts and innervating many glomeruli on the dorso-caudal region. T3 divides itself into at least 3 sub-branches: two running medially (T3a and T3b) and innervating many medial glomeruli, and one running laterally (T3c) innervating caudo-lateral glomeruli. Tract T2 is a much smaller tract that goes from the nerve entrance through the medial part of the lobe neuropil at approximately half depth and innervates only a few medial glomeruli. Tract T4 is another smaller tract, which runs laterally along the outer side of the glomerular region, and innervates a set of tear-shaped glomeruli on the most dorsal part of the antennal lobe, close to the dorsal lobe. Contrary to other glomeruli with a clearly stained cortex, these T4 glomeruli are characterized by a homogeneous staining of sensory neurons. A conspicuous tract of neurons bypasses the antennal lobe completely on its dorso-lateral side and forms the two tracts innervating the dorsal lobe (T5) and the subesophageal zone (T6) that transmit mechanosensory and gustatory information respectively. Single glomeruli from the confocal images were reconstructed (Fig. 1c) and we found 158 ± 4 glomeruli in the antennal lobe of middle-sized bumble bees (n = 4 bumble bees), a slightly lower number compared to honey bees (∼160-166 glomeruli) (Galizia et al., 1999; Kelber et al., 2006; Kirschner et al., 2006), and roughly corresponding to the number of OR genes found in bumble bees (Sadd et al., 2015). We observed that in bumble bees, as in honey bees, the outer surface of the antennal lobe consists of a single layer of glomeruli. This arrangement is particularly well adapted to optical measurements of glomerular activity (see below).

### Anatomy of the bumble bee antennal lobe - projection neuron innervation

Further similarities between honey bee and bumble bee olfactory systems were observed at the level of projection neurons innervation (Strube-Bloss et al., 2015). Using the classical technique used in honey bees (Carcaud et al., 2018; Sachse et al., 1999), we stained the lateral antennal-lobe tract (l-ALT) of projection neurons (Fig. 1a). By introducing tracers into the protocerebrum at a location lateral to the α-lobe of the mushroom bodies and rostral to the lateral horn, we obtained clear staining of l-ALT PNs (Fig. 2a) in ventral glomeruli of the antennal lobe (Fig. 1c, red glomeruli innervated by T1 mainly). In contrast to anterograde staining of OSNs, PN staining was found to be homogeneous in the whole volume of the glomeruli, with PN somata visible on the edge of the glomerular area of the AL (Fig. 2a).

**Figure 2.**
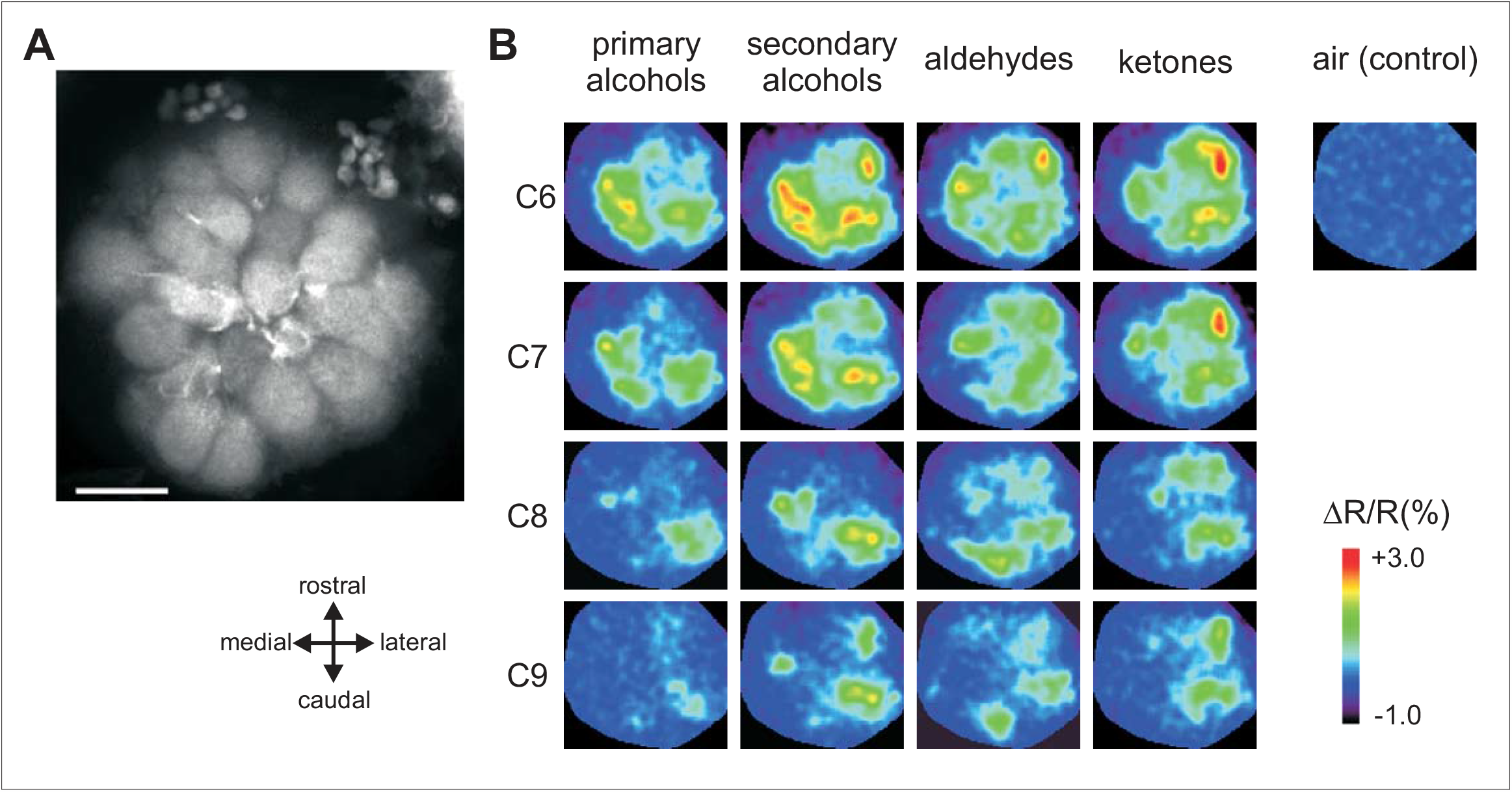
Odor-induced calcium signals from glomeruli innervated by the lateral antennal-lobe tract (l-ALT). **(a)** Confocal image (z-projection over 14 µm, from 6 to 20 µm depth) of the superior part of the AL after retrograde staining (using tetramethylrhodamine dextran) of l-ALT PNs. Dendrites of l-ALT PNs are clearly visible in all observed glomeruli. **(b)** Calcium signals in the AL evoked by a panel of 16 odorants varying systematically according to their carbon chain length (C6–C9) and their functional group (primary and secondary alcohols, aldehydes and ketones). Relative fluorescence changes (ΔR/R%) are presented in a false-color code, from dark blue (no response) to red (maximal response). Different odorants induce different glomerular activity patterns.

All these observations confirm that OSN and l-ALT PN glomerular innervations are highly similar in bumble bees compared to the ones observed in the honey bee olfactory system. We then wondered whether similar olfactory coding rules are found in the two olfactory systems.

### In vivo calcium imaging

We performed *in vivo* calcium imaging measurements in the bumble bee AL using the calcium indicator Fura-2 dextran, and recorded calcium responses from the dendrites of l-ALT PNs in ventral glomeruli (T1 region). We studied the coding of floral odorants. Inspired by previous work on honey bee olfactory perception and coding (Carcaud et al., 2018; Carcaud et al., 2012; Guerrieri et al., 2005; Sachse et al., 1999), we presented to the bumble bees a set of 16 odorants differing systematically in their functional group and chain length.

### Intensity of odor-induced responses

All odorants induced remarkable activity in a combination of AL glomeruli, while air control stimulation had no effect (Fig. 2b). All odorants induced significant activity in comparison to the air control (Fig. 3a; n = 14; RM-ANOVA, F_16, 208_ = 17.1, p<0.0001, comparisons to the control: Dunnett test, p < 0.01). As the odorants systematically varied in terms of chemical group and carbon chain length, we evaluated the effect of these properties on the intensity of calcium responses. Odorants with different functional groups induced different activity levels (Fig. 3b, RM-ANOVA F_3,39_ = 22.1, p < 0.0001). Among functional groups, the weakest responses were evoked by primary alcohols, which induced significantly lower responses than the other chemical groups (Tukey HSD test: p<0.01 compared to secondary alcohols and p<0.001 compared to ketones and aldehydes), which did not differ from each other. Odorants with different chain lengths also induced different activity levels (Fig. 3c, RM-ANOVA, F_3,39_ = 14.4, p < 0.0001). Generally, global response intensity decreased with increasing chain length, i.e. odorant molecules with 6 and 7 carbons induced stronger neural activity than odorants with 8 and 9 carbons (Tukey HSD test: a *vs* b: p<0.01). This pattern of results recapitulates the observations made in honey bees (Carcaud et al., 2018; Carcaud et al., 2012; Sachse et al., 1999) and can be explained by the volatility of the odorants, as measured by their individual vapor pressure. Indeed, AL response was highly correlated to vapor pressure (Fig. 3d, R^2^ = 0.88, F_1,14_ = 106.7, p < 0.0001) confirming that the more volatile the odorant (i.e. the larger its vapor pressure), the more molecules were present in headspace in the sample and the larger was the recorded AL response to this odorant. In the presented odorant panel, alcohols and molecules with longer carbon chains possess lower volatility and thus induced lower responses.

**Figure 3.**
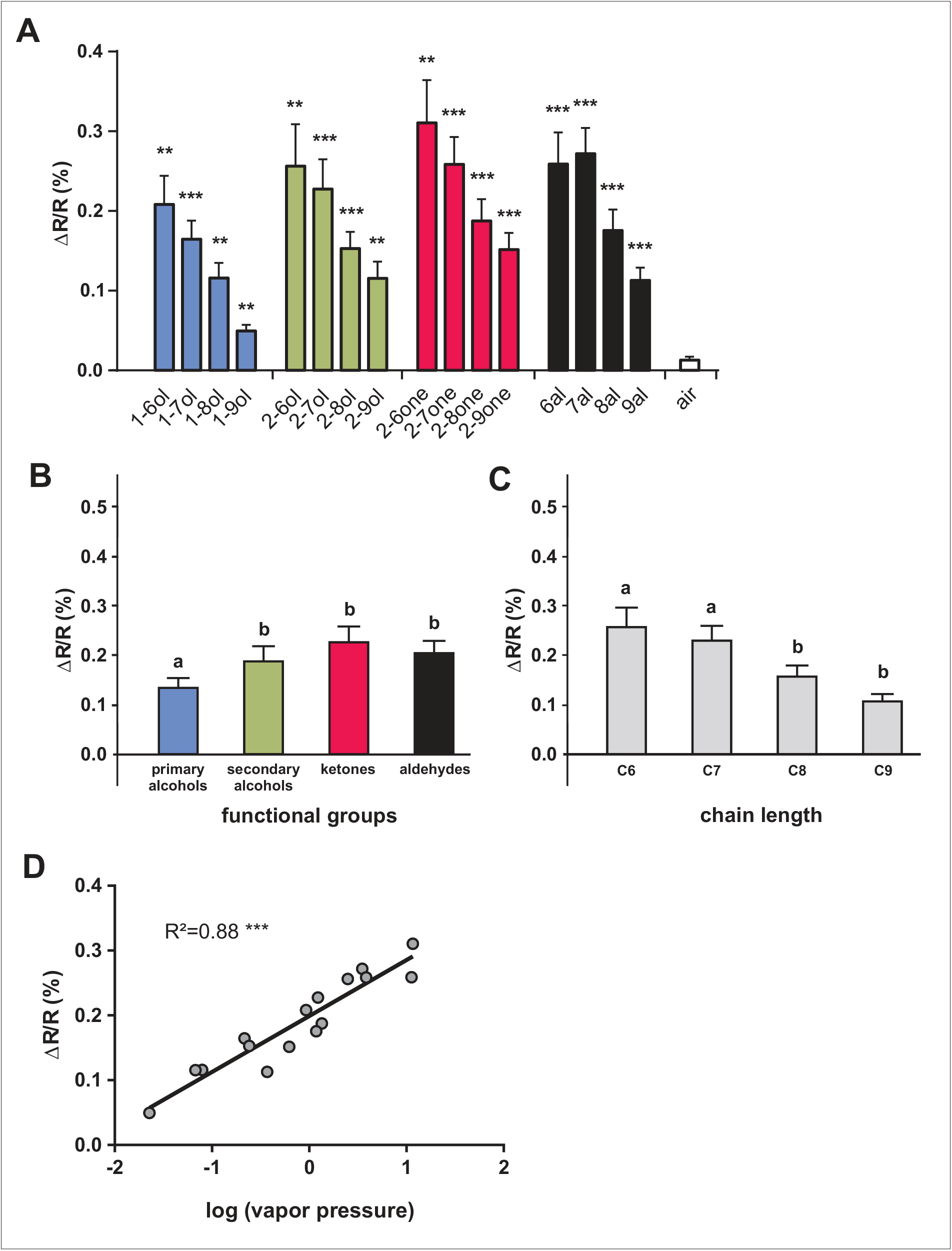
Intensity of calcium responses to 16 aliphatic odorants. **(a)** Amplitude of calcium responses (ΔR/R%) recorded in l-APT PNs to the 16 different odorants. All odors induce significant activity in comparison to the air control (n=14, p < 0.01). **(b)** Amplitude of calcium responses (ΔR/R%) to different odorants according to their functional group (primary and secondary alcohols, aldehydes, and ketones). Primary alcohols (in blue) induced weaker activity than the other functional groups (n = 14, p < 0.01). **(c)** Amplitude of calcium responses (ΔR/R%) depending on odorants’ carbon chain length (6, 7, 8, and 9 carbons). Odorants with the longest carbon chain (C8 and C9) induced weaker activation than odorants with a short carbon chain (C6 and C7) (n=14, p < 0.01). **(d)** Amplitude of calcium responses (ΔR/R%) induced by each of the 16 aliphatic odorants as a function of its vapor pressure (in log units). The linear regression shows a significant correlation (R^2^ = 0.88, p < 0.001).

### Similarity among odor response maps

We then evaluated how chemical characteristics of odorants affected similarity relationships among AL response maps. We thus calculated a measure of (dis-)similarity between response maps (pixelwise Euclidian distance) for all possible pairs of the 16 tested odorants, and produced a distance matrix, which provides an overview of similarity relationships among these odorants (Fig. 4). The more similar odor responses were between two odorants, the smaller are the Euclidian distances and the more intense is the color in the matrix. The matrix reveals a strong effect of the odorant’s carbon chain length on similarity relationships, as shown by the red diagonal lines in the matrix (e.g. for primary alcohols vs. secondary alcohols). Generally, distances between any two odorants of the same carbon chain length were smaller than distances between odorants with different carbon chain lengths. The matrix also suggests that odor pairs with longer carbon chains (C8 vs. C9) evoke more similar activation patterns (i.e. smaller Euclidian distances) than odor pairs with shorter carbon chain length (C6 vs. C7). This more pronounced similarity is also visible in single recordings, as for example shown in Fig. 2b, where a distinct change in the glomerular activity map can be seen between C7 and C8 odorants, but not between C6 and C7 or between C8 and C9 molecules. Odorants’ functional group also plays a role in similarity relationships, although this effect is less easily visible in the matrix. Some pairs of functional groups show higher similarity than others, for instance most primary and secondary alcohols show a high similarity (low distance).

**Figure 4.**
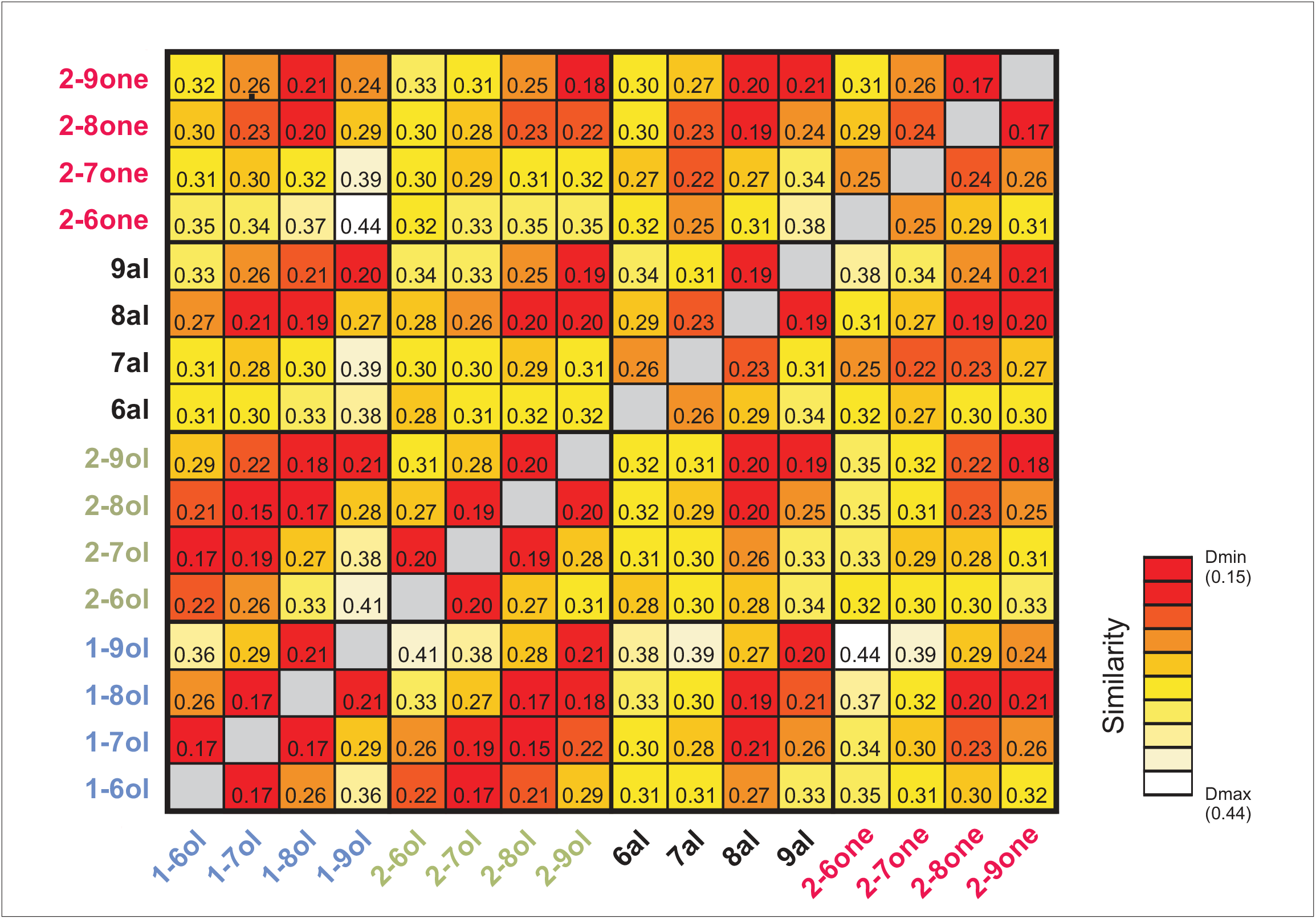
Similarity relationships among the 16 odorants. The matrix shows in a false-color code the Euclidean distances for the 120 odorant pairs. Higher similarity (shorter distances, *D*_min_) is represented in red, while lower similarity (longer distances, *D*_max_) is shown in lighter colors (white and yellow). The matrix shows generally higher similarity (smaller distances) among odorants with a long carbon chain length (C8 and C9, e.g. 8al vs. 2-9one or 9al vs. 1-8ol) compared to the corresponding odor pair combinations within shorter chain lengths, i.e. C6 and C7. High similarity is also observed between primary and secondary alcohols, along a diagonal line showing a dependency on chain length (lower left side of the matrix).

We confirmed these observations by performing multidimensional analyses using these Euclidian distance measures (Fig. 5). A hierarchical cluster analysis using Ward’s classification method (Fig. 5a) showed that the odorants formed three main clusters. Odorants primarily segregated along two branches. The upper branch predominantly grouped odorants with short carbon chain lengths (C6 and C7). Within this branch, odorants were grouped according to their functional groups, with primary and secondary alcohols in one subgroup (C-OH function) and aldehydes and ketones in the other (C=O function). The lower branch exclusively contained odorants with longer carbon chain lengths (C8 and C9). Within this branch, odorants also tended to be distributed according to their functional group, apart from 1-nonanol. This analysis shows, as can be seen in the matrix (Fig. 4) as well as in individual recordings (Fig. 2b), that long-chain molecules evoke highly similar activity patterns, which are less dependent on the functional group than shorter molecules.

**Figure 5.**
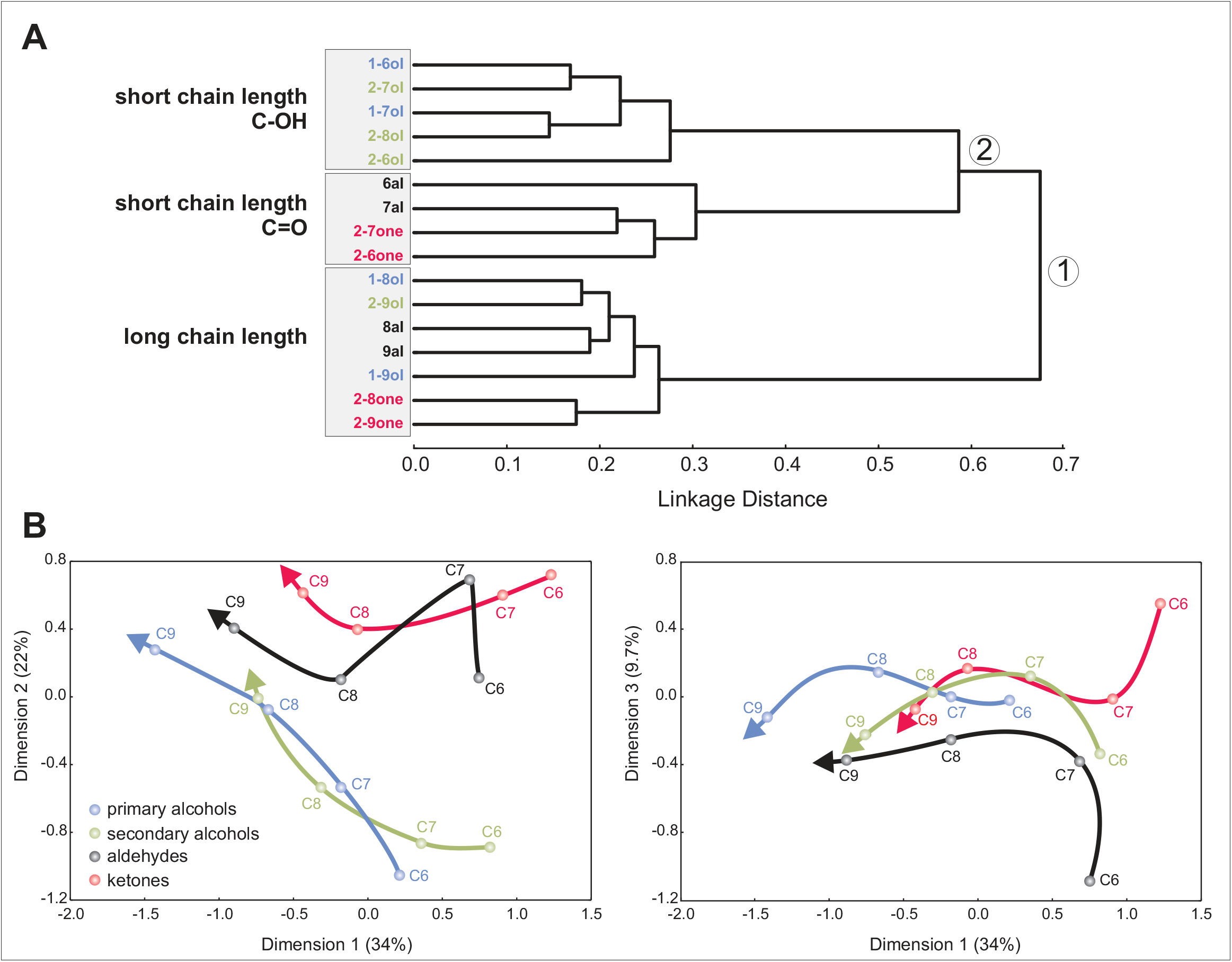
Multidimensional analyses. **(a)** Cluster analysis showing similarity relationships among odorant response maps (Ward’s classification method). Short linkage distance between branches indicates odorants with similar response maps. Functional groups are shown in different colors: primary alcohols in blue, secondary alcohols in green, aldehydes in black, and ketones in red. The analysis shows a first separation (node 1) between odorants with short and long carbon chain lengths. Odorants with a short carbon chain are then subdivided (node 2) into alcohols (primary and secondary, C-OH function) and ketones/aldehydes (C=O function). **(b)** Proximity analysis (multidimensional scaling) based on the Euclidean distance matrix for the 16 odorants. The first dimension (left panel) explains 34% of overall variance and orders molecules according to their chain length from short (on the right, C6 and C7) to long (on the left, C8 and C9). The second dimension (right panel) explains 22.5% of variance and distinctly separates alcohols (blue, green) from ketones (red) and aldehydes (black). Functional group separation is clearer for short-chain than for long-chain molecules. The third dimension (right) explains 9.7% of variance and separates aldehydes (black) from other molecules. Altogether, odorants’ chain length and functional group represent main coding dimensions for odorants in l-ALT projection neurons.

We next performed a proximity analysis (also called multidimensional scaling) on the basis of the Euclidian distance matrix, to understand the most meaningful dimensions underlying similarity relationships among odorants (Fig. 5b). The first three dimensions explained about two thirds of overall variance (dimension 1: 34%; dimension 2: 22%; dimension 3: 9.7%). Dimension 1 mostly provided information about odorants’ carbon chain length, as odorants are represented along this axis by increasing carbon chain length for all functional groups (Fig. 5b left). Dimension 2 contained both functional group and chain length information. Primary and secondary alcohols were not separated from each other, but these functional groups with a C-OH function were clearly separated from both ketones and aldehydes with a C=O function (Fig. 5b left). Dimension 2 also contains carbon chain length information for primary and secondary alcohols, as the odorants are represented along this axis by increasing carbon chain length. Lastly, dimension 3 clearly separates aldehydes (lower values) from ketones, primary and secondary alcohols (higher values, Fig. 5b right). To summarize, the proximity analysis generated three main dimensions, which first represented odorants’ chain length, and then functional group information, distinguishing alcohols, ketones and aldehydes from each other.

The observations made on the distance matrix (Fig. 4) and the multidimensional analyses (Fig. 5) are supported by statistical analyses (Fig. 6). First, odor-specific coding is demonstrated by the fact that odor response maps for presentations of the same odor were more similar (smaller Euclidian distances) than odor response maps for presentations of two different odorants (Fig. 6a, Paired t-test, t = 6.94, p < 0.0001). Second, odorants with the same functional group induced significantly more similar odor response maps compared to odorants with different functional groups (Fig. 6b, Paired t-test, t = 4.69, p < 0.001). Lastly, odorants with the same carbon chain length induced more similar response maps than odorants with different carbon chain lengths (Fig. 6c, Paired t-test, t = 4.99, p < 0.001). This effect increased with the difference in the number of carbon atoms between the odorant molecules (Fig. 6d). The difference between odor maps was thus stronger when the molecules differed by at least 2 carbons, i.e. C6 vs. C8 or C6 vs. C9 (ANOVA F_3,39_ = 25.86, p < 0.0001; Tukey HSD test: a *vs* b p<0.01; a *vs* c p<0.001). These analyses thus demonstrate that odor coding in the bumble bee AL relies on both odorants’ chain length and odorant’s functional group.

**Figure 6.**
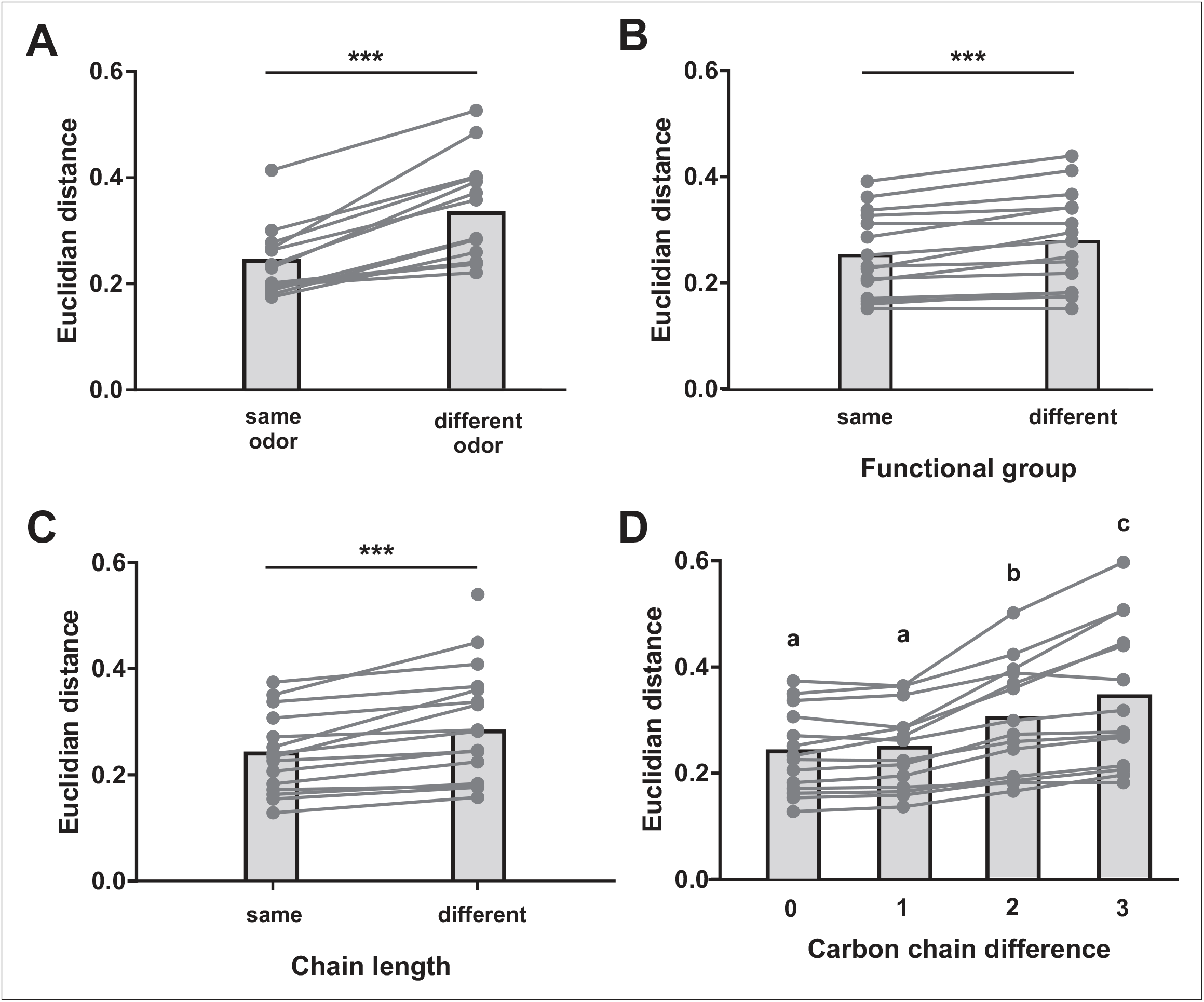
Odor quality coding depending on functional group or carbon chain length information. **(a)** Similarity (Euclidian distance) between presentations of the same or of different odorants. Activity maps are more similar when the same odorant is presented, showing specific odor coding in l-ALT projection neurons (p < 0.001). **(b)** Odorants with the same functional group induce more similar activity patterns than odorants with different functional groups (p < 0.001). **(c)** Odorants with the same chain length induce more similar activity patterns than odorants with different chain lengths (p < 0.001). **(d)** Similarity between odorants depending on the difference in their number of carbon atoms. Euclidean distances increase (i.e. response maps are more dissimilar) with increasing difference in the number of carbon atoms (p < 0.001; a *vs* b, p < 0.01; a *vs* c, p < 0.001; b *vs* c, p < 0.05).

### Comparison of honey bee and bumble bee data

The results we have described so far for bumble bees are generally very similar to the data obtained when imaging the homologous region of the honey bee AL (Carcaud et al., 2018; Sachse et al., 1999). We thus assessed the similarity of odor coding in bumble bees and honey bees by comparing odor-evoked intensity and similarity relationships between the two species. We thus performed linear regression analyses of response intensity (Fig. 7a) and similarity measures between bumble bees and honey bees (Fig. 7b). Response intensities measured for the 16 odorants were highly correlated (R^2^ = 0.57; F_1,14_ = 18.64, p < 0.001) showing that odorants inducing strong responses in bumble bees also induce strong activity in honey bees. Similarly, Euclidian distances between odor response maps for the 120 odor pairs were also strongly correlated between honey bees and bumble bees (R^2^ = 0.55; Mantel test p < 0.0001) indicating that odors inducing similar activity patterns in the honey bee AL also induce similar activity patterns in the bumble bee AL. These comparisons show that odor responses are quite similar between honey bees and bumble bees, both in terms of intensity and similarity relationships.

**Figure 7.**
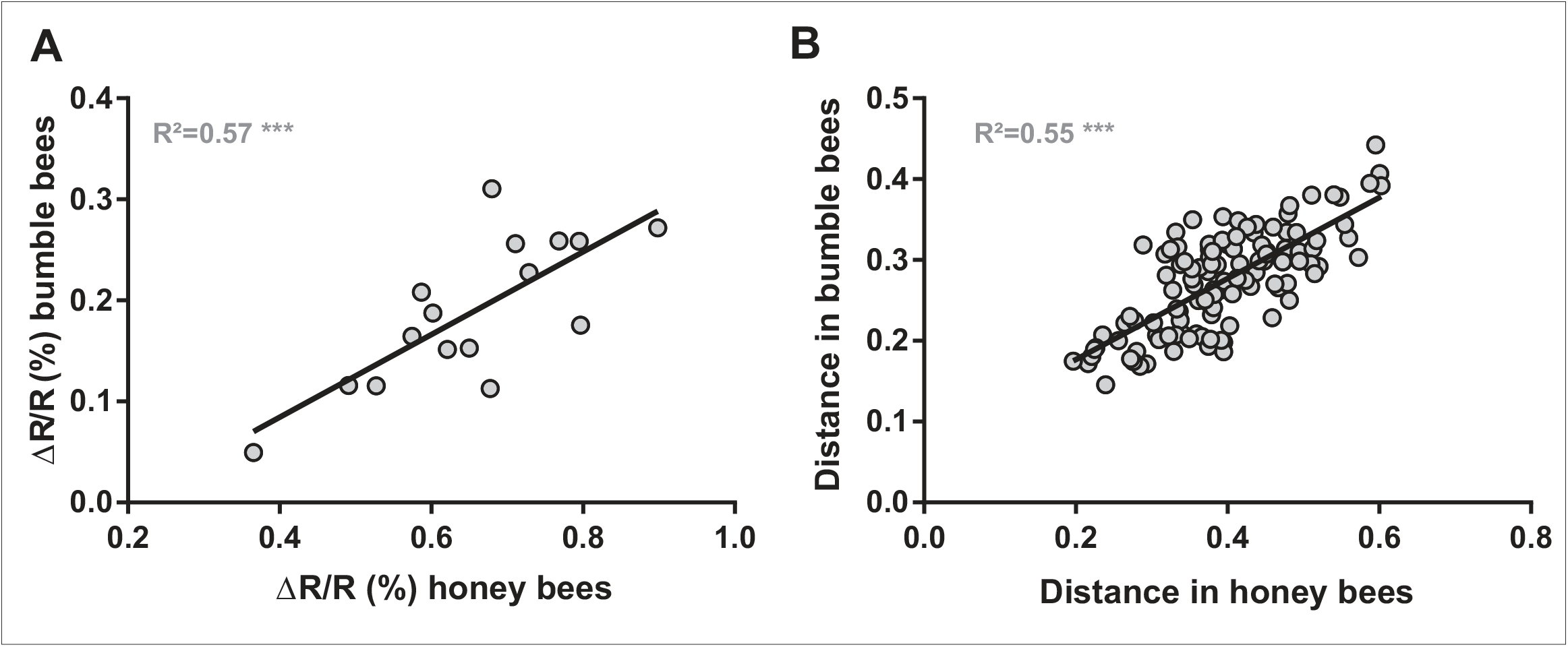
Comparison of odor coding in bumble bee and honey bee AL. **(a)** Correlation of response intensity for each of the 16 presented odorants between bumble bee (n=14) and honey bee (n=10) measures. A high and significant correlation is observed (R^2^ = 0.57, p < 0.001). **(b)** Correlation of Euclidian distances between odor response maps for the 120 odorant pairs obtained in bumble bees and honey bees. A high and significant correlation is also observed (R^2^> = 0.55, p < 0.001). Honey bee data from Carcaud et al., (2018).

## Discussion

This study shows that bumble bees are a suitable model organism for studying central olfactory processing and coding. Using neuroanatomical and neurophysiological approaches, we described AL architecture and measured glomerular activity patterns of in response to odorants. We found that bumble bee l-ALT PNs provide clear information about odorants’ chemical features, here their functional group and chain length. Both the general anatomy of the AL and olfactory coding rules in this structure were found to be closely similar between bumble bees and honey bees.

Neuroanatomical staining and 3D reconstructions indicated that the structure of the bumble bee AL greatly resembles that of the honey bee (Flanagan and Mercer, 1989; Galizia et al., 1999). Both consist of a single layer of glomeruli around an inner coarse neuropil characterized by the presence of numerous local interneurons and projection neurons (see Fig. 1). The restricted innervation of the glomerular cortex by olfactory sensory neurons (OSN) seen in bumble bees is also reminiscent of the honey bee AL (Galizia et al., 1999). To note, in both species, a set of the most dorsal glomeruli innervated by the T4 tract present a homogeneous innervation compared with the exclusively peripheral innervation of the other glomeruli. The existence of the 4 OSN tracts in the bumble bee antennal lobe (Fig. 1b, plus 2 bypassing tracts towards dorsal lobe and subesophageal zone), and their similar arrangement to that observed in the honey bee (Flanagan and Mercer, 1989; Galizia et al., 1999; Kirschner et al., 2006) suggest a strong homology between the olfactory systems of both insects. The most prominent T1 tract innervates a large proportion of glomeruli on the ventral surface of the AL, which are directly accessible when opening the brain capsule. As in honey bees, these glomeruli could be stained retrogradely by placing dye crystals on the l-ALT tract of projection neurons (Fig. 2; Strube-Bloss et al. 2015). We are thus confident that the group of glomeruli that we imaged in bumble bees is structurally homologous to the glomeruli usually imaged in honey bees with the ventral preparation (Joerges et al., 1997; Sachse et al., 1999; Sachse and Galizia, 2002).

The number of glomeruli we have found in bumble bees was slightly lower than the typical number of glomeruli found in honey bees (∼160-165; Flanagan and Mercer, 1989; Galizia et al., 1999). Our values were obtained on medium-sized bumble bee workers. Contrary to honey bees, bumble bees exhibit a pronounced size variation among workers of the same colony (Plowright and Jay 1968). On the one hand, these differences influence the behaviour of the bees, as small bumble bees rather remain within the hive and care for the brood, whereas the largest workers forage more frequently outside of the colony. This size-dependent division of labor is organized as a continuum, so that medium-sized workers care for the brood or manage the temperature of the colony but also go out to forage (Goulson, 2010). On the other hand, body size also influences brain anatomy. Mares and colleagues (Mares et al., 2005) found for instance that the central body and the mushroom body lobes of larger bumble bees are smaller (relative to the whole brain volume) than those of smaller workers. However, it was not the case for the antennal lobe, whose relative size did not change with worker size (Mares et al., 2005). At the periphery, the number of olfactory sensilla, as well as their density, on the antenna were shown to increase with body size in bumble bees (Spaethe et al., 2007). One possibility is that these would express more ORs and consequently display more glomeruli in their AL, as observed in leaf cutting ants, which also show pronounced size-polymorphism (Kelber et al., 2010). In this case, larger workers may show a higher number of glomeruli, reaching the range of glomeruli in the honey bee AL (Flanagan and Mercer, 1989; Galizia et al., 1999). Alternatively, size may not affect glomerulus number, but rather the number of OSNs of each type innervating each glomerulus. The observation that body size increase in bumble bees also correlates with a higher sensitivity towards odorants supports this possibility (Spaethe et al., 2007). Future work should specifically address possible differences in AL organization and glomerulus numbers among differently-sized bumble bee workers.

We retrogradely stained the ventral glomeruli of the bumble bee AL via the lateral projection neuron pathway (l-ALT), which is a common pathway in Hymenoptera (Rössler and Zube, 2011). Using *in vivo* optical recordings, we demonstrated that a panel of 16 aliphatic odorants evokes consistent neuronal activity in the glomeruli of the bumble bee antennal lobe. In particular, we found that an odorant’s functional group and chain length influence the intensity of odor-evoked signals, with primary alcohols inducing significantly lower activity than other functional groups, and short chain molecules activating glomeruli more strongly than molecules with longer chain lengths (Fig. 3). These effects, also found in honey bees (Carcaud et al., 2018; Carcaud et al., 2012; Sachse et al., 1999) are explained by the strong correlation found between response intensity and odorant’s vapor pressure. Thus, the bumble bee antennal lobe, as its honeybee counterpart, does not display any specific sensitivity for any of the odorants in our panel, and AL activity mainly reflects odor concentration in vapor phase.

We then showed that each odorant evokes a specific glomerular activity pattern (Fig. 6a), which is different from that evoked by other odorants. In addition, molecules with a similar chemical structure produced similar activity patterns (see for instance 2-hexanol and 2-heptanol in Fig 1) and olfactory coding was influenced by both chemical features, carbon chain length and functional group. This result was expected as these molecular features strongly influence the chemical similarity among molecules, as measured from 1664 molecular descriptor values (Haddad et al., 2008). Similarly, carbon chain length and functional group were shown to affect neural coding both in invertebrates (Carcaud et al., 2018; Carcaud et al., 2012; Couto et al., 2005; Dupuy et al., 2010; Sachse et al., 1999) and in mammals (Johnson & Leon, 2007; Mori et al., 2006). Accordingly, these molecular features influence olfactory perception in humans (Beets, 1970; Laska & Teubner, 1999; Nagao et al., 2002; Polak, 1973) and honey bees (Guerrieri et al., 2005; Laska & Teubner, 1999).

Odor-evoked activity within PNs is the product of OSN activity entering the AL and of local inhibitory networks carrying out local computations (Carcaud et al., 2018; Sachse & Galizia, 2002, 2003). Despite a highly similar organization of their olfactory pathways, the main difference between honey bee and bumble bee systems lies in the repertoire of ORs expressed at the periphery. A recent study analyzed the genomes of 2 Bombus species, *Bombus terrestris* and *Bombus impatiens* (Sadd et al., 2015) and aiming to identify key players in the evolution of sociality, they compared the genomes of these species with that of the honey bee *Apis mellifera*. Concerning chemoreception, they found that *Bombus* genomes contain a slightly less diverse OR family than *Apis mellifera*, with 159 intact OR genes (excluding 5 pseudogenes). The number of glomeruli that we found in the AL of *Bombus terrestris* in our reconstructions (n=158 ± 4) corresponds well to the number of OR proteins found in the genome of this species, fitting with the general hypothesis in insects that each OSN expresses one type of odor-specific receptor, while all OSNs carrying the same receptor project to the same glomerulus in the AL (Vosshall et al., 2000). This hypothesis is especially appealing as in honey bees the number of olfactory receptor genes largely coincides with the number of glomeruli in the AL (∼165 glomeruli and ∼163 intact OR genes excluding pseudogenes; Robertson and Wanner, 2006). Note however, that in *Drosophila melanogaster*, several AL glomeruli are not innervated by OSNs expressing OR family genes, but rather by neurons expressing ionotropic receptors (Grabe et al., 2016; Silbering et al., 2011). As bumble bee and honey bee genomes each contain about ∼20 IR genes, with several orthologous genes, a proportion of their ALs may be innervated by IR expressing sensory neurons.

The comparison of the honey bee and the bumble bee OR family genes also showed duplications of genes in one or both species, several large species-specific gene lineage expansions, and at least 22 gene losses, reflecting the birth-and-death gene family evolution typical of these receptors (Ramdya & Benton, 2010). Recent evidence in different species of the genus *Drosophila* suggests that the number of olfactory receptor genes has remained quite similar for the entire period of *Drosophila* evolution (63 million years; Tamura et al., 2004), but that frequent gains and losses of genes occurred in each evolutionary lineage (Nozawa and Nei, 2007). This may have changed the sequence of olfactory receptor neurons leading to different glomerular wiring patterns. The oldest common ancestors of honey bees and bumble bees are estimated to have lived between 70 and 90 million years ago (Michener and Grimaldi, 1988; Schultz et al., 2001; Ramírez et al., 2010). This long time of separate evolution suggests that profound changes could also have taken place in the sequence of olfactory receptor genes in both species, which may have led to a complete change of each receptor’s sensitivity spectrum to odorant molecules, as well as their localization in the AL. In accordance with these observations, when we compared the patterns evoked by the different odorants in the two species, we found it very difficult to identify homologous glomeruli. Our finding of highly similar olfactory coding *rules*, supported by a clear coding of chain length and functional group information, in bumble bees as in honey bees should not be construed as a contradiction to the points raised above. In our experience, this finding is a by-product of the joined sensitivities and selectivities of the numerous OSN/ORs imaged simultaneously. In a previous study, we showed that similarity relationships among inter-odorant maps measured in the ALs of an ant (*Camponotus fellah*) and the honey bee using calcium imaging were similar to those measured in the rat olfactory bulb using an utterly different recording technique (2-deoxyglucose autoradiography) (Dupuy et al., 2010). The general rule was simple: odorants with a similar molecular structure (chain length and/or functional group) induced similar activity patterns in each insect’s antennal lobe as well as in this mammal’s olfactory bulb. More generally, it was observed that similarity relationships in a range of different species, including invertebrates and rodents, could be predicted to some extent based on purely molecular descriptors of odorants (Haddad et al., 2008). Thus, an emerging property of the multiple coding channels of each system (the ORs) which each detect a different but overlapping range of odorant molecule features is that neural representations mirror the chemical characteristics of the molecules. Importantly, this emerging property does not depend on the type of receptors expressed at the periphery since it is well established that olfactory receptor (OR) proteins in insects and vertebrates are unrelated (Benton, 2015; Benton et al., 2006). Coming back to our study, honeybee and bumble bee ORs may well have evolved independently for a long time, our study shows that the two neural ensembles that were recorded on the ventral surface of their ALs perform a reliable depiction of odorants’ chemical features, granting these insects with a clear representation of odorants’ structure. In honey bees, we previously showed that inter-odorant similarity relationships in the AL could predict bees’ behavioral responses in a generalization protocol, so that similar odorants in the AL were treated as similar by the bees in their behavior (Guerrieri et al., 2005; Carcaud et al. 2012, 2018). The high correlation we found between odor-evoked response maps in the bumble bee and honeybee ALs suggests that bumble bees are indeed equipped, like honey bees, to respond to odorants according to chemical dimensions. We thus predict that future behavioral experiments in bumble bees shall reveal a similar organization of their olfactory perceptual space based on odorants chemical dimensions, as found in honey bees (Guerrieri et al., 2005).

In conclusion, our study unravels a high similarity in the general organization of the primary olfactory processing center of bumble bees and honey bees. In addition, it shows similar olfactory coding rules conveying each system with a reliable depiction of odorants’ chemical structure. While we concentrated here on the coding of general odorant features, we expect that future studies devoted to the coding of species-specific odorants, like social pheromones, may reveal more remarkable differences between both systems.

## Methods

### Bumble bee preparation

Medium-sized bumble bee workers were caught from an indoor colony (Koppert, Berkel en Rodenrijs, The Netherlands) and chilled on ice for 5 min until they stopped moving. Then, bumble bees were prepared following the standard preparation used to image the AL in honey bees (Carcaud et al., 2018; Joerges et al., 1997; Sachse & Galizia, 2002). In summary, the bumble bee’s head was inserted and fixed in a plastic chamber with its antennae oriented to the front of the chamber. Using beeswax, the proboscis was flued at the front end of the holder to avoid movement of the brain during the experiment. Hairs on the top of the bumble bee head were removed and a pool was built with beeswax and pieces of plastic around the rostral part of the head capsule (behind the antennae). The pool was made waterproof with two-component epoxy glue (red Araldite, Bostik Findley, S.A.). A small window was then cut in the head cuticle from the bases of the antennae up to the ocelli, and glands as well as parts of the tracheal sheath were removed to expose the antennal lobes and parts of the protocerebrum. Finally, the pool was filled with some ringer solution (in mM: NaCl, 130; KCl, 6; MgCl_2_, 4; CaCl_2_, 5; sucrose, 160; glucose, 25; Hepes, 10; pH 6.7, 500 mOsmol; all chemicals from Sigma-Aldrich, Lyon, France), to avoid desiccation of the brain surface. Three hours prior to the experiment, a dye mixture was inserted into the brain with a broken borosilicate micropipette, aiming for the tract of l-ALT projection neurons, between the α lobe and the border of the optic lobe, rostrally from the lateral horn. The dye mixture consisted of the calcium-indicator Fura-2 dextran (10,000 kDa, Life technologies, France) and of tetramethylrhodamine dextran (10,000 kDa, Life technologies, France) for later anatomical observation, both in bovine serum albumin (2%).

### Calcium imaging

*In vivo* optical recordings were performed as described elsewhere (Carcaud et al., 2018; Mota et al., 2013; Roussel et al., 2014), with a T.I.L.L. Photonics imaging system (Martinsried, Germany), under an epifluorescence microscope (Olympus BX51WI) with a 10× water-immersion objective (Olympus, UMPlanFL; NA 0.3), which was dipped into the ringer solution covering the brain. Only one AL was recorded in each bumble bee. Images were taken with a 640 × 480 pixels 12-bit monochrome CCD camera (T.I.L.L. Imago) cooled to −12°C. Fura-2 was alternatively excited with 340 nm and 380 nm monochromatic light (T.I.L.L. Polychrom IV). Each measurement thus consisted of 50 double frames recorded at a rate of 5 Hz (integration time for each frame at 340 nm: 40–80 ms; for 380 nm: 10-20 ms) with 4 × 4 binning on chip (pixel image size corresponded to 4.8 µm × 4.8 µm). The filter set on the microscope contained a 490 nm dichroic filter and a bandpass (50 nm) 525 nm emission filter.

### Odor presentation

A constant clean airstream, into which odor stimuli could be presented, was directed from a distance of 2 cm to the bumble bee’s antennae. Odor stimuli (see below) were given at the 15^th^ frame for 1 s (5 frames). Odor sources consisted in exchangeable Pasteur pipettes containing a piece of filter paper (1 cm^2^) soaked with 5µl of pure odorant (Sigma Aldrich, France).

In a first experiment, we tested 16 different aliphatic odorants that are part of floral blends bumble bees encounter while foraging (Knudsen et al., 1993). The odorants differed systematically in terms of their carbon chain lengths (between 6 and 9 carbon atoms) and their functional groups (primary alcohol, secondary alcohol, aldehyde and ketone). As control stimulus, we used a pipette containing a clean piece of filter paper without odor solution. This stimulus set was also used in a recent calcium imaging study of PN responses in the honey bee AL (Carcaud et al., 2018) allowing the comparison of odor coding in honey bees and bumble bees. The olfactory stimuli were presented three times in a pseudo-randomized order, avoiding consecutive stimuli to contain the same functional group or the same carbon chain length.

### Data processing and analyses

Data were analyzed using custom-made software written in IDL 6.0 (Research Systems, Boulder, CO) (Carcaud et al., 2015). Each odor presentation produced a four-dimensional array consisting of the excitation wavelength (340 or 380 nm), two spatial dimensions (x- and y-coordinates) along time (50 frames). First, the fluorescence ratio between excitation wavelengths at each pixel and time point was calculated: R = F_340nm_ / F_380nm_. The relative fluorescence changes were then computed between the recorded odor responses R at each time point compared to the background fluorescence (before any odor presentation) R_0_, defined as the average of the three images before odor stimulus onset (frames 12-14). Relative fluorescence changes were thus calculated as: ΔR = (R – R_0_) / R_0_. The two spatial dimensions were then filtered with a gaussian filter of window size 7×7 pixels to reduce photon noise. Lastly, possible irregularities of lamp illumination were corrected by subtracting the median pixel value of each frame from each single pixel of the corresponding frame. The amplitude of the odor-induced response was calculated by subtracting the average of three consecutive frames during the odor presentation (frames 17-19) from the average of 3 frames before stimulus onset (frames 12-14).

Activity maps (Fig. 2) represent the average amplitude observed over the three presentations of each odorant, in a false-color code, from dark blue (no signal) to red (maximum signal). As unambiguous identification of identical glomeruli across individual bumble bees was not feasible, odor coding was analyzed over the entire surface of the AL using a pixelwise analysis that avoids any bias due to glomerular misidentification. It was previously shown in honey bees that results based on the pixelwise method lead exactly to the same conclusions as glomerular identification (Carcaud et al., 2015). For each bee, a mask was precisely drawn along the edges of the AL to limit the measure of odor-evoked responses to the glomerular area. Global glomerular activity upon odor stimulation was measured by averaging the intensity values of all pixels within the unmasked area. Evaluation of (dis-)similarity relationships between odorant representations was performed by calculating pixelwise Euclidian distances for all pairs of the 16 odorant stimuli used (120 odor pairs). For all analyses, average values for the three presentations of each odorant were used except for the comparison of Euclidian distances for the *same or different* odorants (Fig. 6a), where each single odorant presentation was used.

### Anatomical staining

For antennal staining of the whole antennal lobe, the scapes of the antennae were carefully opened using a microscalpel, and the antennal nerve was cut with a borosilicate micropipette coated with tetramethylrhodamine dextran (10,000 kDa, Life technologies, France). Afterwards, animals were kept in a cool place until the next day to allow the dye to migrate to the AL and to stain OSN processes within the glomeruli. The brains were removed and fixed in 4% paraformaldehyde solution for at least 24 h. They were then dehydrated in ascending concentrations of ethanol, cleared and stored in methyl salicylate (Sigma-Aldrich, Lyon, France). Images of the tetramethylrhodamine-stained glomeruli were taken using a confocal laser scanning microscope (Zeiss, LSM 700) with a W Plan-Apochromat 20x/1.0 objective and a 555 nm excitation wavelength at 2 µm optical section thickness and pixel size of 0.31 µm × 0.31 µm. Recorded stacks of images were adjusted in brightness and contrast using imageJ (Rasband; National Institutes of Health, Bethesda, MD). Segmentation and anatomical reconstruction of the antennal lobe was performed using Amira (version 4.5.1 Mercury Computer Systems, Merignac, France).

After successful calcium imaging, the brains were removed and the same techniques as described above were used.

### Statistical analysis

Normality of the data was tested and confirmed for almost all data points using Shapiro-Wilk normality test. We thus applied parametric statistics over the whole study. When normality was not achieved for all data points in an analysis, the corresponding non-paramatric test was performed. In all cases, both types of tests gave the same result, and therefore the text only describes parametric results. The intensities of responses to the different odorants were compared using ANOVA for repeated measurements. When significant, Dunnett’s test was applied to compare the intensity of each response to a common reference, the air control. Odor-evoked response intensities between functional groups and chain lengths were compared using ANOVA for repeated measurements, followed by Tukey post-hoc tests for further analysis of statistically significant main effects. Paired t-tests were applied to compare Euclidian distances obtained for different presentations of the same odor *vs* presentations of different odors, as well as for odors with the same or with a different functional group or chain length. A Pearson correlation analysis was performed between response intensity and the logarithm of odorants’ vapor pressure. In some analyses (Fig. 7), data recorded in bumble bees were compared to data recorded in honey bees (Carcaud et al., 2018), using exactly the same experimental and analytical procedures. A Pearson correlation analysis thus evaluated a possible correlation of odor-response intensities between the two species. A Mantel test was used to evaluate a possible correlation between bumble bee and honey Euclidian distance matrices. All tests were performed with GraphPad Prism (version 7, GraphPad software) or R (www.r-project.org). All values are displayed as means ± SEM.

## Acknowledgements

The study was supported by the Deutsche Forschungsgemeinschaft (DFG). We thank Pr. Martin Egelhaaf for providing additional financial support for M.M. We are grateful to Maud Combe for developing the custom programs used for data analysis. We thank the ANR (Projects,2010-BLAN-1712-01 and ANR-17-CE20-003 to J.C.S.) and the French Research Ministry (J.C.). We also would like to thank Antoine Couto and Manon Lefèvre Houard for their help with the anatomical reconstruction.

## Authors contributions

A preliminary version of this work appeared in the doctoral thesis of the first author (Universität Bielefeld, 2013). M.M., J.C., M.E. and J.C.S. conceived the experiments. M.M. collected the data. M.M., J.C. and J.C.S. analyzed the data and interpreted the results. M.M, J.C. and J.C.S. wrote the manuscript. All authors read and approved the final version of the manuscript.

## Data availability statement

All data are available upon request from the corresponding author.

